# A proteogenomic resource enabling integrated analysis of *Listeria* genotype-proteotype-phenotype relationships

**DOI:** 10.1101/668053

**Authors:** Adithi R. Varadarajan, Maria P. Pavlou, Sandra Goetze, Virginie Grosboillot, Yang Shen, Martin J. Loessner, Christian H. Ahrens, Bernd Wollscheid

## Abstract

*Listeria monocytogenes* is an opportunistic foodborne pathogen responsible for listeriosis, a potentially fatal foodborne disease. Many different *Listeria* strains and serotypes exist, but a proteogenomic resource that bridges the gap in our molecular understanding of the relationships between the *Listeria* genotypes and phenotypes via proteotypes is still missing. Here we devised a next-generation proteogenomics strategy that enables the community to rapidly proteotype *Listeria* strains and relate this information back to the genotype. Based on sequencing and *de novo* assembly of the two most commonly used *Listeria* model strains, EGD-e and ScottA, we established two comprehensive *Listeria* proteogenomic databases. A genome comparison established core- and strain-specific genes potentially responsible for virulence differences. Next, we established a DIA/SWATH-based proteotyping strategy, including a new and robust sample preparation workflow, that enables the reproducible, sensitive, and relative quantitative measurement of *Listeria* proteotypes. This reusable and publically available DIA/SWATH library covers 70% of open reading frames of *Listeria* and represents the most extensive spectral library for *Listeria* proteotype analysis to date. We used these two new resources to investigate the *Listeria* proteotype in states mimicking the upper gastrointestinal passage. Exposure of *Listeria* to bile salts at 37 °C, which simulates conditions encountered in the duodenum, showed significant proteotype perturbations including an increase of FlaA, the structural protein of flagella. Given that *Listeria* is known to lose its flagella above 30 °C, this was an unexpected finding. The formation of flagella, which might have implications on infectivity, was validated by parallel reaction monitoring and light and scanning electron microscopy. *flaA* transcript levels were not significantly different with and without exposure to bile salts at 37 °C, suggesting regulation at the post-transcriptional level. Together, these analyses provide a comprehensive proteogenomic resource and toolbox for the *Listeria* community enabling the analysis of *Listeria* genotype-proteotype-phenotype relationships.

## Introduction

*Listeria monocytogenes* is a highly adaptable environmental bacterium that can exist both as plant saprophyte and as animal pathogen [1]. The Gram-positive, rod-shaped, facultative anaerobic bacterium is the causative agent of listeriosis [2,3]. Although the incidence of listeriosis is relatively low compared to other common foodborne diseases, it is associated with one of the highest mortality rates (20% – 30%) [4]. *Listeria* strains are categorized into at least 14 serotypes [5], among these three (1/2a, 1/2b, and 4b) are responsible for the majority of clinical cases. EGD-e, a widely used model system of serotype 1/2a, is the serotype most frequently recovered from foods or food-processing plants. In contrast, ScottA is a widely used model system of serotype 4b, which causes the majority of human epidemics [5]. Infection by *L. monocytogenes* usually occurs after digestion of contaminated foods and in individuals with impaired cell-mediated immunity. The elderly, immunosuppressed patients, pregnant women, and neonates are particularly susceptible [2]. An infection may lead to meningitis, sepsis, or, by crossing the placenta, infection of the fetus and subsequent abortion [5]. Upon ingestion, *L. monocytogenes* must resist multiple stresses encountered in the gastrointestinal (GI) tract, including variation in pH, osmolarity, and bile salts [6,7]. Notably, survival in the GI tract is a prerequisite to establish a successful infection in the host [8].

*L. monocytogenes* has served as a key bacterial model system to study host pathogen interaction [9], and studies of *L. monocytogenes* have led to the discovery of several new concepts in biology [10]. These included the discovery of unconventional mechanisms regulating bacterial gene expression, including the first RNA thermosensor regulating virulence [11], the excludon concept [12], and the discovery of an atypical member of the CRISPR family devoid of *cas* genes [10]. Additionally, research on *L. monocytogenes* has contributed to a better understanding of the structure and dynamics of the host cell cytoskeleton with the discovery of the first actin nucleator in eukaryotic cells (the Arp2/3 complex) [13,14] and the elucidation of a novel role for clathrin in actin polymerization [10]. Finally, analysis of *L. monocytogenes* has been instrumental in characterizing naïve-to-memory CD8 T cell generation and differentiation [15].

The number of ‘omics datasets that study different aspects of *L. monocytogenes* biology has increased exponentially in recent years. These datasets include a growing number of Illumina-based fragmented genomes [16] as well as complete genome sequences [17], the latter of which provide an optimal basis for comparative genomics and functional genomics studies. Such comparisons have helped to identify *Listeria* virulence factors and regions associated with pathogenicity [18,19]. Moreover, analyses of complete genomes enabled detailed investigations of genes transcribed under clinically relevant conditions [12] and enabled identification of novel protein coding genes and the correct protein N-termini/start sites through proteogenomics [20]. Yet, changes in transcript abundances upon perturbation often do not correlate with abundance changes of the corresponding protein products [21,22], and transcriptional data alone does not provide important functional information about post-transcriptional regulation such as protein modifications [23] or changes in protein interaction networks or cell-surface remodeling of the host following an infection [24]. Such information can, however, be obtained using state-of-the-art proteotype profiling.

The proteotype is defined as the state of the proteome at a particular time [25]. A proteome therefore consists of many proteotypes. The proteotype concept takes the dynamic nature of the proteome into account and extends it to the organization of proteins and their co-existing proteoforms in time and space [25]. Mass spectrometry-based proteotype profiling has matured recently through technological and methodological advances allowing for increased depth of proteome coverage [26–29] and sample throughput. This launches the next generation (next-gen) proteomics era of comprehensive and quantitative proteotype profiling. Recognizing the unmet need to integrate these datasets and to enable meta-analysis in a user-friendly manner, pioneers in the *Listeria* field developed the interactive Listeriomics website (https://listeriomics.pasteur.fr). At the time of publication, it contained 83 *Listeria* genome, 492 transcriptome, and 74 proteome datasets [30]. Notably, the majority of proteomic studies (qualitative and quantitative) were based on 2D gel studies, so modest numbers of proteins have been quantified thus far [31–37]. Similarly, the workflows employed to date have not provided the depth required for quantitative systems-level characterizations at the protein level.

In the present study, we set out to generate and validate two proteogenomic resources to enable analysis of genotype-proteotype-phenotype relationships in *L. monocytogenes* strains. By relying on the *de novo* assembled genomes of the model strains ScottA and EGD-e, we generated a mass spectrometry-based toolbox using next-gen DIA/SWATH workflows to enable the sensitive, repetitive, and quantitative interrogation of *L. monocytogenes* proteotypes. DIA/SWATH-MS (for Data Independent Acquisition/Sequential Window Acquisition of all THeoretical Mass Spectra) is a recently introduced proteotyping technology based on mass spectrometry, which, compared to data-dependent acquisition (DDA) MS strategies, enables more sensitive generation of comprehensive proteotype datasets. The dynamic range of DIA/SWATH-MS is in excess of four orders of magnitude [38], which matches the dynamic range of the proteotypes expected for our *L. monocytogenes* strains. Recently, proteogenomic studies on *Streptococcus pyogenes* [39] and *Mycobacterium tuberculosis* [40] utilizing the DIA/SWATH technology have illustrated its impact, which led to new biological insights with respect to invasiveness of clinical isolates and to dormancy and resuscitation, respectively.

We applied our newly developed proteogenomic resources and toolbox to investigate how *L. monocytogenes* cells cope with stress encountered during passage through the upper GI tract, a prerequisite for systemic infection of the host. We uncovered evidence for the unexpected expression of flagella upon exposure to bile salts at 37 °C, a condition mimicking the duodenum. The comprehensive proteogenomic resource and toolbox we established here will enable further analyses of the *Listeria* genotype-proteotype-phenotype relationships.

## Results & Discussion

### A generic, genomics-driven strategy enabling the investigation of genotype-proteotype-phenotype relationships

We selected two widely used *L. monocytogenes* strains, EGD-e and ScottA, as model systems to evaluate the applicability of our genomics-driven next-gen proteomics workflow (Fig 1). EGD-e and ScottA belong to serotypes 1/2a and 4b, respectively, which are responsible for the majority of listeriosis cases (S1 Table); ScottA is more invasive than EGD-e [41]. We relied on an integrated workflow to obtain a quantitative profile of the *Listeria* proteotype. This workflow contains four main components: genomics, comparative genomics, proteotype analysis, and proteogenomics (Fig 1).

**Fig 1.**
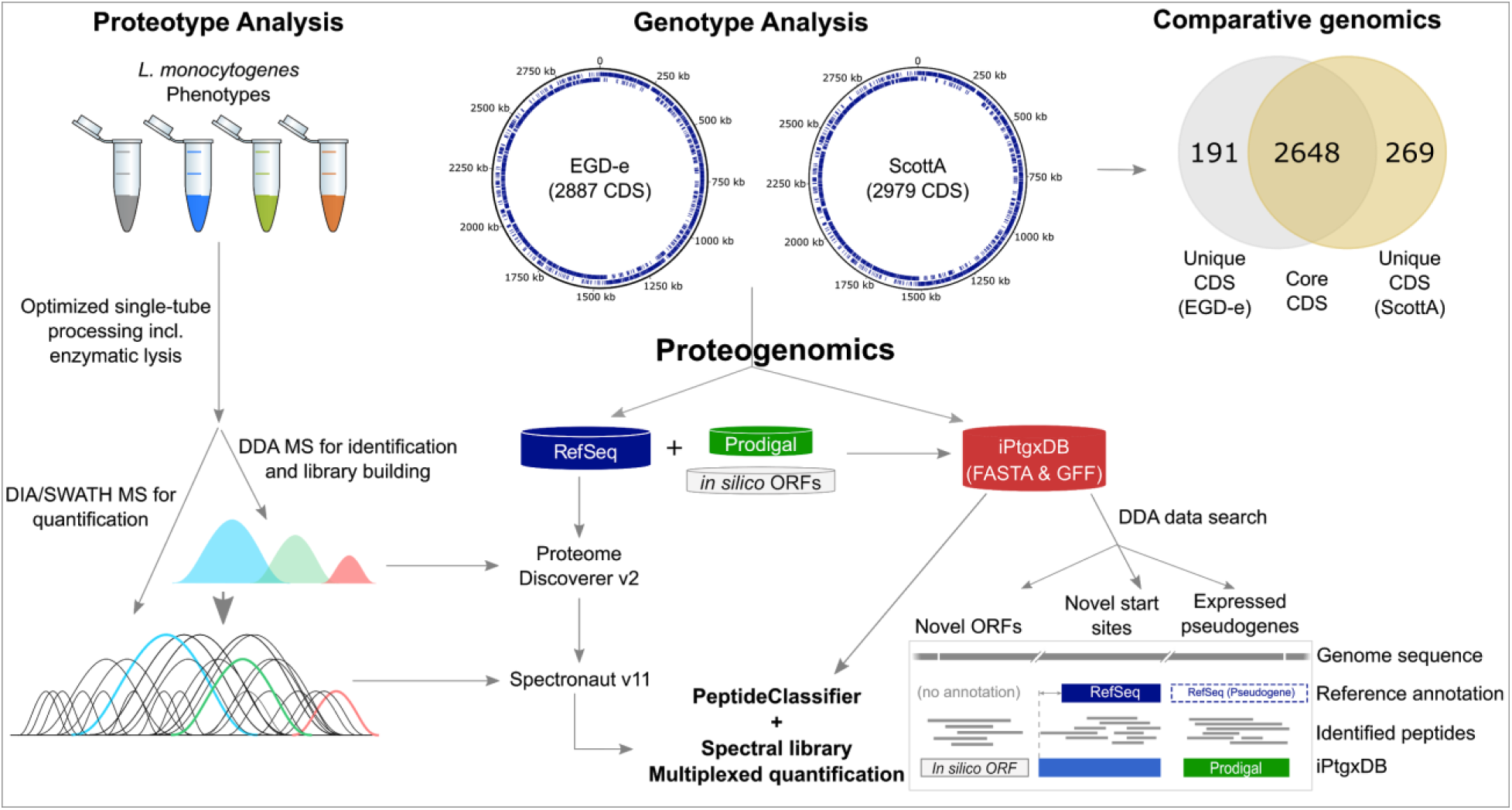
Overview of our next-gen proteogenomics workflow for *L. monocytogenes*. A sample preparation method was developed that allows rapid, reproducible single-tube reactions for lysis, digestion, desalting, and subsequent proteotype profiling with DDA- and DIA-MS (left panel). In parallel, the genomes of both EGD-e and ScottA were *de novo* assembled into complete, high-quality genome sequences, and their RefSeq annotations were obtained from the NCBI’s prokaryotic genome annotation pipeline (PGAP) [42] (middle upper panel). Comparative genomics identified both shared core gene clusters and gene clusters unique to EGD-e and ScottA (right upper panel). Moreover, an *ab initio* gene prediction based on Prodigal and an advanced *in silico* (six-frame translation) annotation (see Materials and Methods) were integrated with the RefSeq annotation to obtain a minimally redundant iPtgxDB [43] for EGD-e and for ScottA. DDA-based proteomics data were searched against the publicly available iPtgxDBs (https://iptgxdb.expasy.org) to obtain proteogenomic evidence for novel open reading frames (ORFs), novel start sites, and expressed pseudogenes (right lower panel). For the proteotype analysis, proteomics data obtained from DDA mode were searched against the RefSeq annotations and spectral libraries were generated with Spectronaut. These publicly available resources were then used to analyze and quantify proteins obtained from DIA mode.

The first step in our workflow was a genomics analysis. Although a complete NCBI reference genome sequence existed for EGD-e, the NCBI reference genome sequence for ScottA was incomplete and consisted of five contigs [44]. Motivated by our recent finding of significant differences between an NCBI reference genome and the *de novo* assembly of the actual lab strain [43], and an earlier study on *Pseudomonas aeruginosa* PAO1 that had demonstrated substantial genomic fluidity between closely related strains [45], we sequenced and *de novo* assembled both genomes to create the best possible reference sequence for the two strains, an important aspect for the proteogenomics element. Next, a comparative genomics analysis was carried out to identify core and strain-specific genes, which, upon integration with protein abundance data, might provide clues to explain the different phenotypes of the strains. Thirdly, we performed DDA-MS experiments under relevant conditions to obtain extensive *Listeria* proteotype datasets, which became the basis for the construction of *Listeria* spectral libraries. After establishing a rapid and reproducible sample preparation workflow, we were able to quickly and quantitatively profile *Listeria* under various conditions and perturbations by DIA/SWATH. Lastly, in addition to the standard proteotype search against generic protein databases such as NCBI RefSeq and Uniprot, a search was carried out against a specialized, integrated proteogenomics search database (iPtgxDB; see Materials and Methods) [43], allowing us to identify protein expression evidence for as of yet unannotated small ORFs (smORFs), additional N-terminal start sites, and expressed pseudogenes (Fig 1).

A variant of such a proteogenomics approach recently revealed N-terminal peptides both from internal start sites of annotated *L. monocytogenes* EGD-e proteins and from six novel, unannotated smORFs including Prli42 [20]. This 31-amino acid protein relays oxidative stress signals to the stressosome to activate the general stress-sensing pathway, the sigma B regulon, and represents the long-sought link between stress and the stressosome [46]. Despite the many important functions of smORFs, such small, functional open reading frames are often missed in current genome annotations [47]. Proteogenomics and ribosome profiling have emerged as the most important technologies for comprehensive identification of smORFs [47]. Consequently, we added a proteogenomics element to our generic strategy. This has two benefits: First, proteogenomics can be included in the initial genome annotation, thereby increasing its quality [43,48]. A public website (https://iptgxdb.expasy.org/) supports this for newly sequenced genomes [43], such as isolates from microbiomes or type strains from the Genomic Encyclopedia of Bacteria and Archaea (GEBA) initiative, which aims to expand the phylogenetic diversity of completely sequenced prokaryotic genomes [49]. Second, the identification of more comprehensive protein catalogs including functionally relevant smORFs, will better enable studies to model systems based on quantitative data of all functional elements.

### Complete genome sequences of EGD-e and ScottA and comparative genomics

An analysis of the repeat complexity of all publicly available, completely sequenced genomes of *L. monocytogenes* strains (status: March, 2018) revealed that almost 95% of the roughly 150 strains are so-called “class I” genomes, which are straightforward to assemble (few repeats, none longer than the rDNA operons of up to 7 kb) [17]. In contrast, eight strains also had few repeats overall, but those present were up to 11 kb in length [17]. Using long-read Pacific Biosciences (PacBio) sequencing data (including a BluePippin size selection step; see Materials and Methods) and the assembly algorithm HGAP3 [50], we were able to *de novo* assemble one complete chromosome for EGD-e (2.94 Mbp) and one for ScottA (3.03 Mbp) with PacBio read coverages of 260x and 280x, respectively (Fig 2). To correct remaining homopolymer errors and to remove small insertions or deletions (INDELs) in the PacBio data [51], both strains were also sequenced using the highly accurate short-read Illumina protocol. The Illumina data allowed us to ensure that no additional plasmids could be assembled that might have been lost in the BluePippin size selection step before creating the insert libraries for PacBio sequencing. Additional genome properties, including the number of protein-coding genes predicted by NCBI’s PGAP annotation pipeline, are listed in S2 Table. Two intact and one putative prophage were identified by PHASTER [52] in ScottA, and two putative prophages were identified in EGD-e (Materials and Methods).

**Fig 2.**
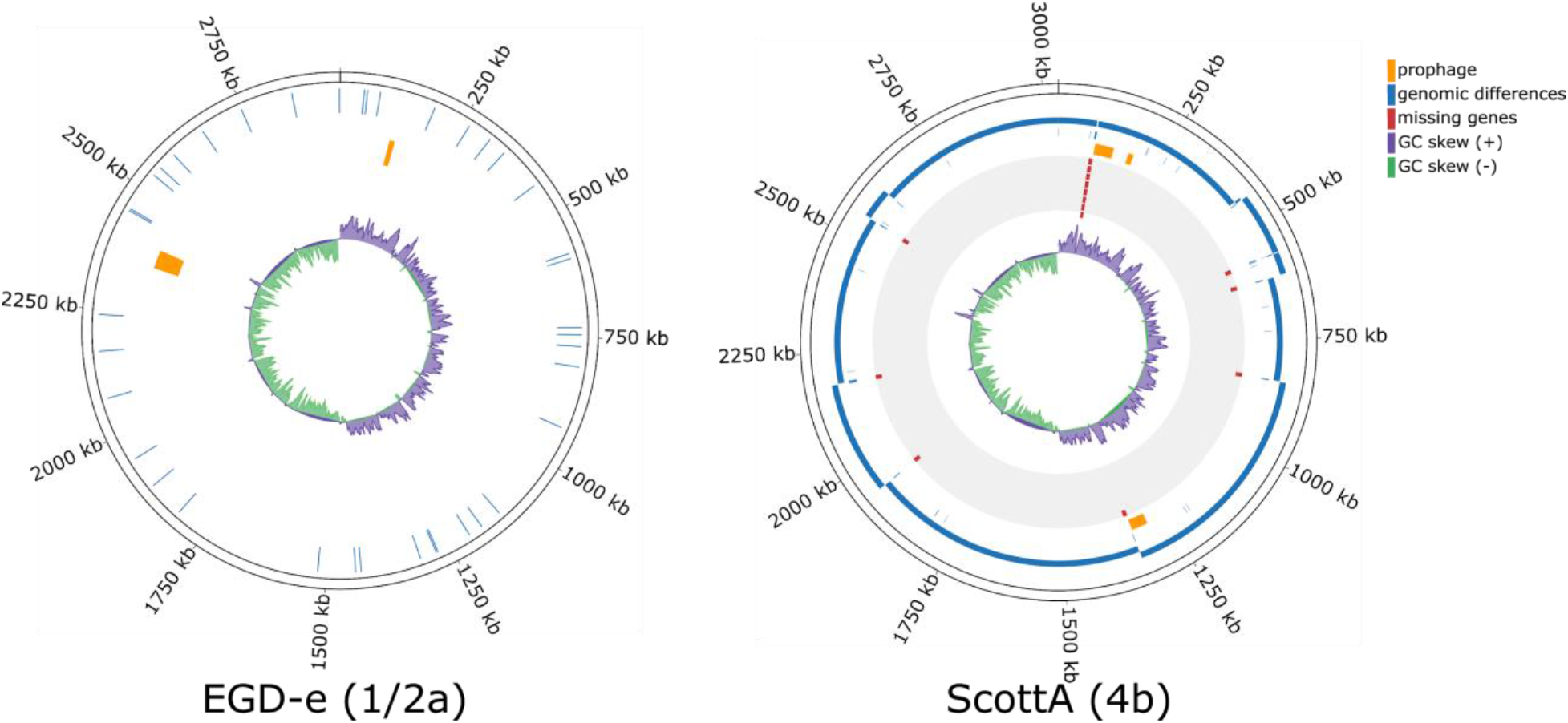
Circular plots showing the *de novo*-assembled genome sequences (outer ring with genomic coordinates) of EGD-e and ScottA and the genomic differences compared to the existing reference sequences, NC_003210 and NZ_CM001159, respectively. Whereas our EGD-e assembly exhibited minor differences compared to the NCBI reference genome (SNVs and INDELS in blue, second circle), there were a number of differences between our complete ScottA assembly and the NCBI reference (incomplete, 5 contigs), which had been assembled using a reference-based assembly approach [44]. The mapping of the five contigs and the remaining gaps (blue) are shown in the second circle; prophages (orange) in the third, 15 missing genes (red) in the fourth, and the GC skew (positive, purple; negative, green) in the fifth circle.

Comparing our *de novo* assembled ScottA genome to the NCBI reference (CM001159.1; 5 contigs), we observed a total of 11,953 base pairs (bp) of missing sequence, which affected 14 genes that were completely or partially missed in the earlier, at the time state of the art, reference-based genome assembly [44]. The missing genes included seven hypothetical proteins, three surface proteins, one cell-wall anchor-domain containing protein, and three transposases (S3 Table). The genomic differences included 14 insertions, 16 deletions, 93 single-nucleotide variations (SNVs), and 34 variations affecting two or more nucleotides (Fig 2). In contrast, only 28 SNVs and 11 single-bp INDELs were observed between our EGD-e assembly and the NCBI reference genome (NC_003210.1) (Fig 2).

The example of ScottA illustrates that a *de novo* assembly strategy is preferable over a reference-based one, which can easily miss important genome sequence differences. Roughly 570 of 9,331 bacterial genome assemblies (6.1%) that we recently analyzed [17] were prepared using a reference-based genome assembly strategy; these assemblies thus have to be treated with caution. With current long-read sequencing technologies like Pacific Biosciences and Oxford Nanopore Technologies readily generating sequence reads longer than 20 kb, *de novo* genome assembly has become the preferred approach.

Finally, complete genomes also represent the best basis for comparative genomics studies, as core genes have been missed when more fragmented assemblies based on short Illumina reads were used for comparative genomics [53]. A comparison of the complete genomes of EGD-e and ScottA using Roary [54] revealed a high similarity with 2,648 core gene clusters (orthologous proteins), 191 (6.6%) EGD-e-specific and 269 (9%) ScottA-specific gene clusters (Fig 1 and Table 1). Notably, five of the fourteen genes that were missed in the five contigs of the fragmented ScottA genome are indeed core genes that are shared by both strains. A list of core and strain-specific genes is provided in S3 Table. These results will enable the *Listeria* research community to further explore differing virulence capabilities and other properties of these two strains. To facilitate such comparisons and integration with other data sets, we also provide a detailed table with the genes of both strains, functional annotations, proteomic abundance evidence, and a reciprocal best BLAST hit comparison against the ListiList EGD-e proteins with identifiers in the form of LmoXXXX (https://listeriomics.pasteur.fr), where X is a number from 0-9) in S4 Table. Together, these new proteogenomic resources represent a high-quality set of puzzle pieces required for modeling and understanding the *Listeria* life cycle and infectious mode of action.

**Table 1.** Overview of the comparative genomics results of the newly sequenced and assembled EGD-e and ScottA genomes including core and strain-specific protein-coding genes.

|  | *L. monocytogenes*<br>EGD-e | *L. monocytogenes*<br>ScottA |
| --- | --- | --- |
| Genbank accession # | CP023861 | CP023862 |
| Size of the chromosome (bp) | 2,944,523 | 3,030,813 |
| Total number of protein-coding genes | 2,887 | 2,979 |
| Number of strain-specific genes | 191 | 269 |
| Number of core genes | 2,648 | 2,648 |

### Growth under conditions mimicking passage through the gastrointestinal tract

Despite intense research, knowledge of the proteotype adaptations that facilitate survival of *L. monocytogenes* in the GI tract remains incomplete. To cause a systemic infection, the bacteria have to be ingested and travel through the GI tract [8]. The GI tract is a hostile environment, and the bacteria are subjected to mechanical and chemical stresses that differ depending on the compartment: These include acidity (stomach), exposure to bile (duodenum), and exposure to high osmolarity (jejunum) (Fig 3A). To enable a systems-wide, quantitative characterization of the proteins required for survival and adaptation of *L. monocytogenes* under these GI-imposed stresses, we first sought to devise *in vitro* culturing conditions that resemble the different microenvironments of the GI tract. As not all genes will be expressed under one condition, these perturbations were chosen in order to obtain a broad representation of expressed *Listeria* proteins. We reasoned that discovery-driven DDA-based proteotype data from such conditions would allow us to create a comprehensive spectral library, an important element of our next-gen DIA/SWATH-based proteotyping strategy (Fig 1). This was achieved by culturing *L. monocytogenes* cells in three different conditions at 37 °C [55] and in a control condition (buffered peptone water (BPW), pH 7.4) in which the cells did not replicate (based on OD_600_ measurements, data not shown).

**Fig 3.**
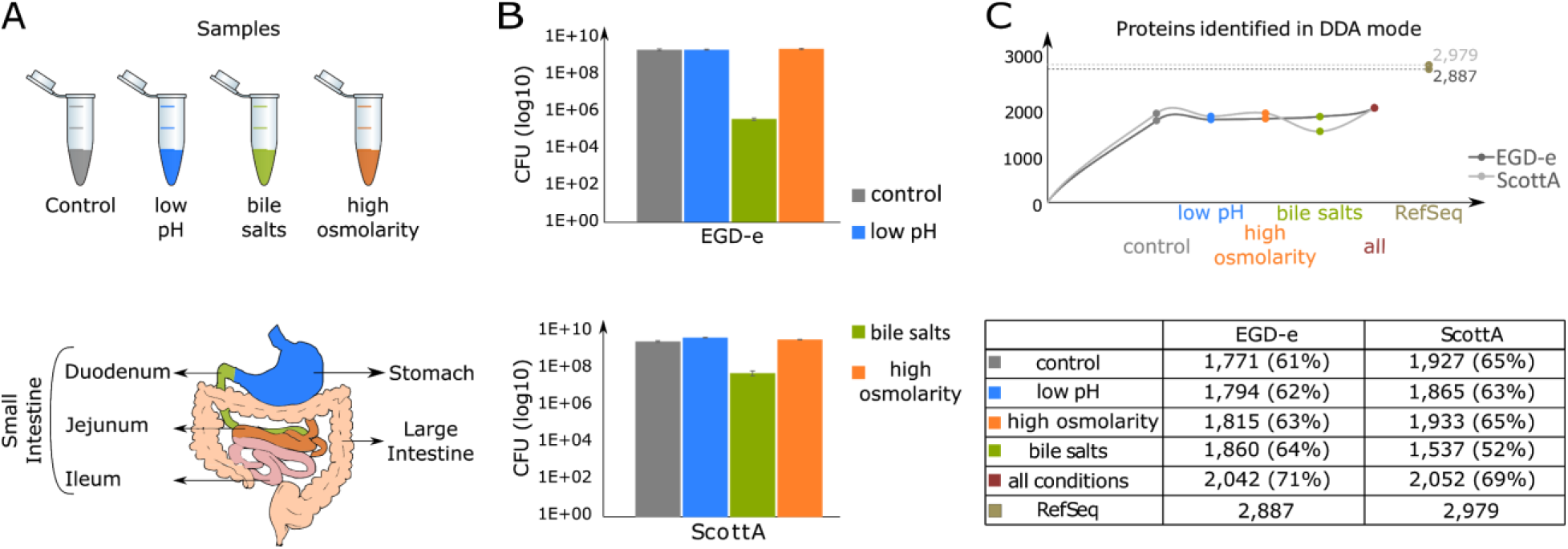
Discovery-driven DDA-based *Listeria* protein expression data obtained under conditions mimicking passage through the upper GI tract. **(A)** Sample conditions (control, grey; stomach, blue; duodenum, green; jejunum, orange). **(B)** Viability of cells in control and three different conditions. **(C)** Number of proteins identified per condition for each strain, the overall number of identified proteins (red), and the number of proteins annotated in each strain (brown).

After testing several combinations (data not shown) the following buffer compositions were chosen (Fig 3A): To mimic conditions encountered in the stomach, the pH of the buffer was 4, and the medium included 1,000 units/ml of pepsin. To simulate the duodenum, the medium included 0.3% w/v bile salts (pH 7.4). Finally, in order to mimic conditions encountered in the jejunum, the pH of the medium was 8, and it contained 0.3 M sucrose to increase the osmolarity (Fig 3A). The viability of the strains was not affected by the stresses of low pH/pepsin and high pH/high osmolarity (Fig 3B). In contrast, viability was severely compromised by the presence of bile salts, indicating that detergent-like activities are detrimental to survival of bacterial cells. In the presence of bile salts, the viability of ScottA (approximately one log lower than in the control) was affected much less than that of EGD-e (approximately 4 logs lower compared to the control) (Fig 3B). The greater sensitivity of EGD-e to both pH and bile compared to other *L. monocytogenes* strains, including ScottA, has been noted before [6,7]. Although these are clearly model conditions, in the absence of better *in vivo* models, they will lead to a better understanding of the mechanisms by which *Listeria* adapts to the host GI tract, informing the development of novel treatment or prophylactic strategies.

### *Listeria* proteotype analysis using a single-tube workflow and spectral libraries

To ensure reproducible, sensitive, and quantitative proteotype measurements both for this study and as a general resource for the *Listeria* research community, a robust sample preparation workflow was needed that included a minimal number of sample preparation steps, while simultaneously enabling a comprehensive protein identification and quantification across conditions/proteotypes. Typically, bacterial cells are lysed mechanically (i.e., by bead beating) or by use of detergents. However, these methods require additional steps for bead or detergent removal making the workflows more tedious, increasing technical variability, and potentially leading to loss of low abundance proteins. Therefore, an efficient and proteotype analysis-compatible workflow was developed in which all steps from lysis to protein digestion are performed in a single tube (Fig 1). Effective, rapid, and complete cell lysis was achieved by incubating *L. monocytogenes* cells with the bacteriophage endolysin (Ply511; see Materials and Methods). Recombinantly produced endolysins applied exogenously to susceptible bacteria display the same lytic properties as their native counterparts [56]. In combination with indirect sonication, combined LysC/trypsin protein cleavage into peptides and desalting using a mixed cationic exchange resin, the sample processing strategy enables rapid proteotyping of *L. monocytogenes*.

To generate spectral libraries, samples were first analyzed in DDA-MS mode. For this, the raw files were analyzed with MaxQuant “dependent peptide” settings [57] to identify the most prominent post-translational modifications, which were then used as variable modifications in a second search, thereby limiting the overall search space. In total, we identified qualitative expression data for between 1,700 and 1,800 proteins per condition for EGD-e and between 1,500 and 1,900 proteins per condition for ScottA (Fig 3C), similar to previously reported, extensive MuDPIT datasets from *Listeria* [36,58]. We next used the discovery-driven DDA-based data to generate spectral libraries for DIA/SWATH-based acquisition and reliable protein quantification across conditions. Summaries of the spectral library characteristics for the two strains are depicted in Table 2 and Fig 4.

**Table 2.**
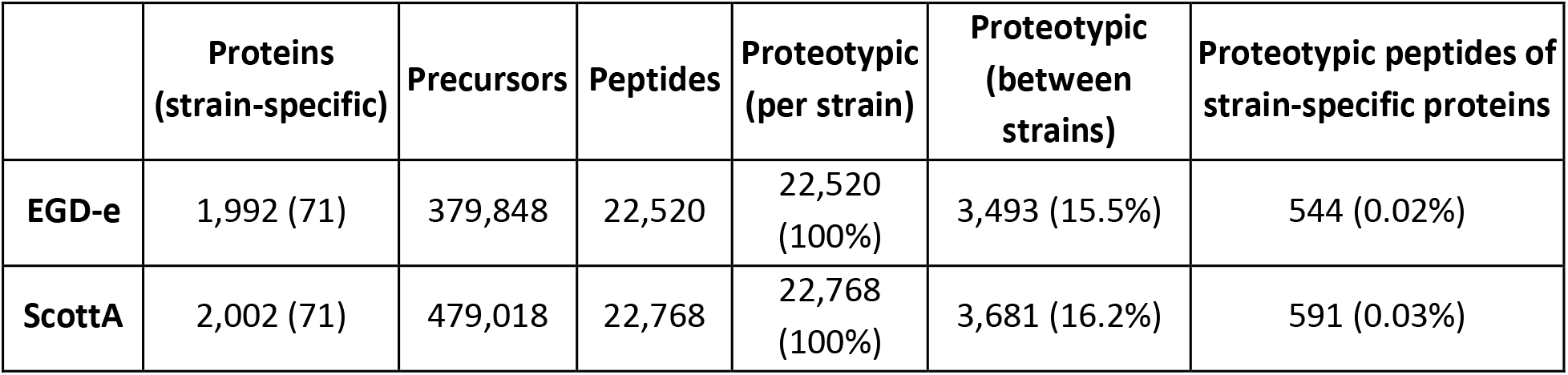
The numbers of precursors, peptides, and proteins that are covered in the spectral libraries of *L. monocytogenes* strains EGD-e and ScottA and peptides that were observed for strain-specific proteins.

**Fig 4.**
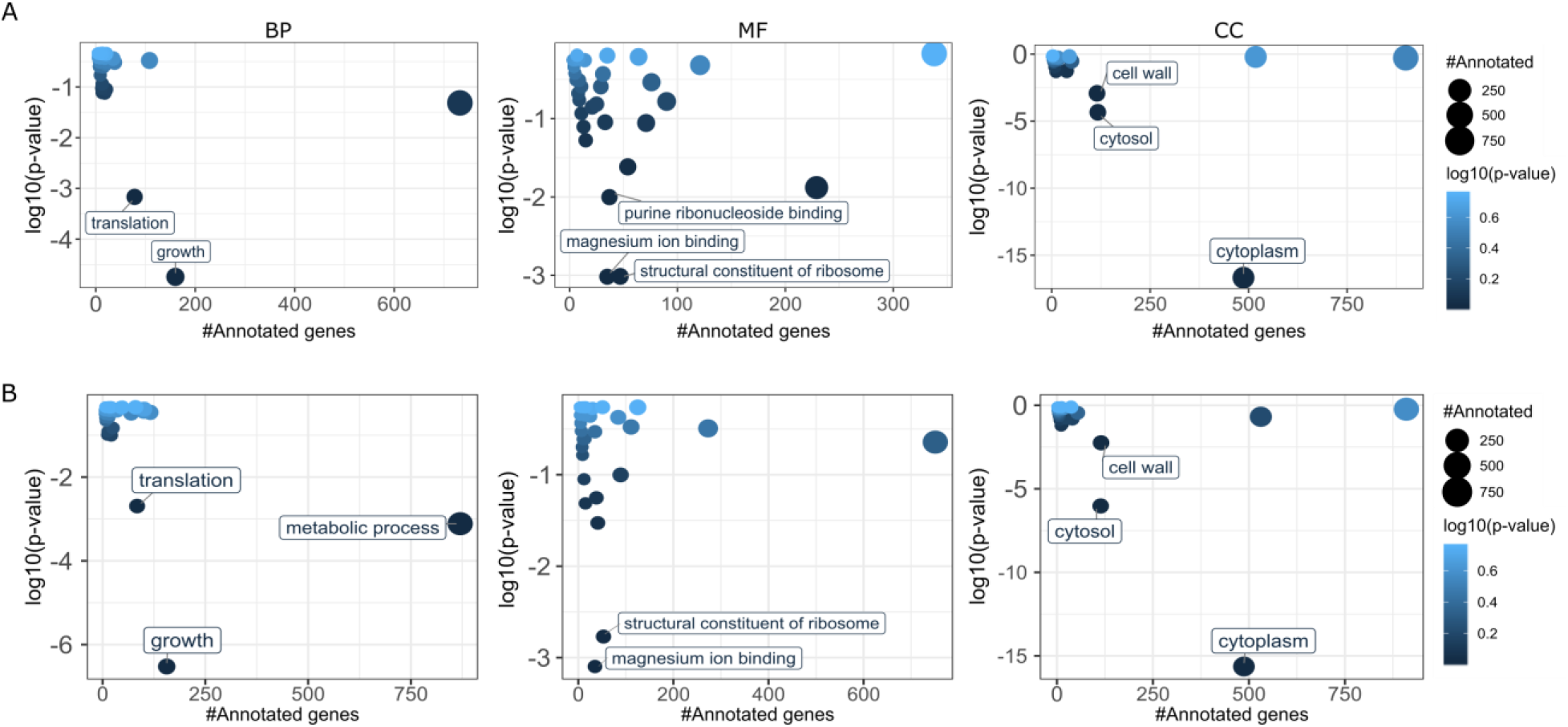
GO enrichment analysis of proteins included in the spectral libraries. GO categories across the three domains biological process (BP), molecular function (MF), and cellular component (CC) were analyzed with a Fisher’s exact test (p-value = 0.01). The X-axis shows the number of annotated genes in each GO category, and the Y axis shows the log10 p-value. Significantly enriched GO categories (p-value < 0.01) in the spectral library are labeled in the figure. The GO category “cytoplasm” had the most significant e-value. **(A)** Plots for strain EGD-e. **(B)** Plots for strain ScottA.

Each library contained roughly 22,000 peptides that corresponded to approximately 2,000 proteins (false discovery rate (FDR) < 1%)). All of the peptides were proteotypic for the given strain as indicated by a PeptideClassifier analysis [59] when using the peptide and protein identifications returned from ProteomeDicoverer as input (see Materials and Methods). Overall, the spectral libraries contained roughly 69% and 67% of the annotated ORFs of EGD-e and ScottA, respectively. To our knowledge, this is the first report of *Listeria-specific* spectral libraries covering such a high proportion of the theoretical proteome. A Gene Ontology (GO) enrichment analysis of the proteins contained in the spectral libraries showed that the majority of pathways were represented in both of the libraries and that similar GO categories were enriched in both strains across the three domains. Notably, the cellular component (CC) ‘cytoplasm’ was significantly over-represented among proteins in the spectral libraries of both strains (Fig 4), whereas proteins associated with the CC category ‘transmembrane’ were under-represented (data not shown). This is an expected outcome and can be explained by our focus on the development of a single-tube, rapid and reproducible sample processing workflow. Proteome coverage without a bias against the membrane proteome has only very rarely been achieved and requires elaborate strategies (computational and biochemical and/or subcellular fractionation) and substantial effort [27]. The CC categories ‘cytosol’ and ‘cell wall’ were also over-represented.

The peptides from the two libraries were mapped against the protein-coding sequences of the other genome to identify strain-specific proteotypic peptides and to assess whether some of the strain-specific protein-coding genes were expressed at the protein level. This mapping was done using an in-house tool that extends the original version of PeptideClassifier [59] thereby enabling proteogenomics for prokaryotes [43]. Only 15.5% and 16.2% of peptides from EGD-e and ScottA, respectively, were strain-specific proteotypic, indicating that most peptides could be used to quantify protein abundance in both strains and potentially also in other *Listeria* strains. Additionally, we found that 544 unambiguous peptides of the EGD-e library confirmed expression of 71 of the 191 EGD-e specific protein-coding genes and that 591 Scott-A strain-specific proteotypic peptides in the library confirmed expression of 71 of 269 ScottA-specific proteins (Table 2 and S3 Table). These efforts resulted in a second new proteogenomic resource that enables fast proteotyping of *L. monocytogenes* using a single-tube processing strategy in combination with a DIA/SWATH-MS-based workflow that benefits from the generated *L. monocytogenes* spectral libraries. These libraries are now publicly available. Due to the high sensitivity of this spectral library-based DIA/SWATH MS approach in combination with top-end MS instruments, it is now conceivable to gain proteotype information directly from limited numbers of cells extracted from *in vivo* models of *L. monocytogenes* infection.

### Integrated proteogenomic databases as a basis to relate proteotype data back to the genotype

In order to make the generated information accessible for public use, we also created an iPtgxDB for each strain. The concept of iPtgxDBs as a “one-stop shop” for a protein search database that combines the benefits of manual curation efforts with the ability to identify missed smORFs by capturing the entire protein-coding potential of a prokaryotic genome has been described previously [43]. Proteotype data from any condition can be searched against the FASTA file, and experimental evidence (peptides or gene expression data, if available) can be integrated with the GFF file provided, thereby allowing users to visualize experimental evidence for novelties (Fig 1). Alternatively, users may simply compare different annotation resources of a genome sequence or even NCBI RefSeq releases, which can differ substantially.

The minimally redundant iPtgxDBs for *L. monocytogenes* EGD-e and ScottA strains contained 65,393, and 67,150 proteins, respectively, and were created by integrating and consolidating annotations from RefSeq [42], Prodigal [60], and a modified form of a six-frame translation using the public iPtgxDB web server (https://iptgxdb.expasy.org/iptgxdb/submit/) [43]. Metadata on the number of proteins in each annotation source, the progressive increase of annotation clusters, and the overall number of ORFs in the final iPtgxDB are shown in S5 Table. Proteotype data measured in the control and three GI-mimicking conditions in DDA-MS mode were searched individually against the iPtgxDB fasta file of each strain (see Materials and Methods).

At a stringent peptide-spectrum match (PSM) level FDR (0.05%, resulting in a protein-level FDR well below 1%), we obtained unambiguous peptide evidence for 1,907 proteins in EGD-e including 1,899 RefSeq proteins and eight additional novel proteins (S6 Table). The novel proteins included two Prodigal-predicted proteins or proteoforms, five *in silico* ORFs, and one protein with an alternative start site. Furthermore, we observed peptide evidence supporting seven of 28 SNVs in our *de novo* assembly compared to the EGD-e reference (NC_003210). In ScottA, we observed unambiguous peptide evidence for 1,910 RefSeq proteins including four proteins (3 transposases and 1 cell wall protein) that were missed in the incomplete ScottA reference sequence (Genbank accession: NZ_CM01159; 5 contigs). Additionally, we identified evidence for six novel proteins including three Prodigal proteins, three *in silico* ORFs, and two alternate protein start sites (S6 Table). Identified novelties include a 45-aa hypothetical protein predicted only by Prodigal that was identified in strain EGD-e by two peptides and 44 PSMs (Fig 5A). Notably, the same 45-aa protein was also identified with these two peptides and 35 PSMs in strain ScottA (data not shown). A BLAST search showed that this smORF is conserved across *L. monocytogenes* strains. The second example is the case of a pseudogene predicted by RefSeq in EGD-e (520 amino acids), which contains an internal, C-terminal stop codon. Protein expression evidence was observed for a corresponding smaller Prodigal-predicted protein of 426 amino acids (Fig 5B). Both the Prodigal and Refseq proteins are annotated as formate-tetrahydrofolate ligase. Peptide evidence confirmed expression of the protein to the internal stop codon; this proteoform also contains the P-loop nucleoside triphosphate hydrolase domain that is conserved across *L. monocytogenes*. The third example shows yet another Prodigal prediction for a protein that is 10-aa longer than the corresponding RefSeq protein (which is 212 amino acids) of strain ScottAin (Fig 5C); a peptide supporting the longer proteoform was identified with 26 PSMs. The same peptide was also identified in strain EGD-e with 25 PSMs (data not shown). Finally, one peptide (EVAEELGVHES<u>A</u>VSR, 11 PSMs) supports the non-synonymous amino acid change (caused by an SNV that results in a threonine to alanine codon change) in the protein annotated in our *de novo* assembly as RNA polymerase factor sigma-54 of 447 amino acids (Fig 5D). S7 Fig shows proteomics evidence (3 peptides, 17 PSMs) for another Prodigal-predicted protein (207 amino acids) annotated as a RpiR family phosphosugar-binding transcriptional regulator; the corresponding Refseq annotation wrongly predicts a pseudogene.

**Fig 5.**
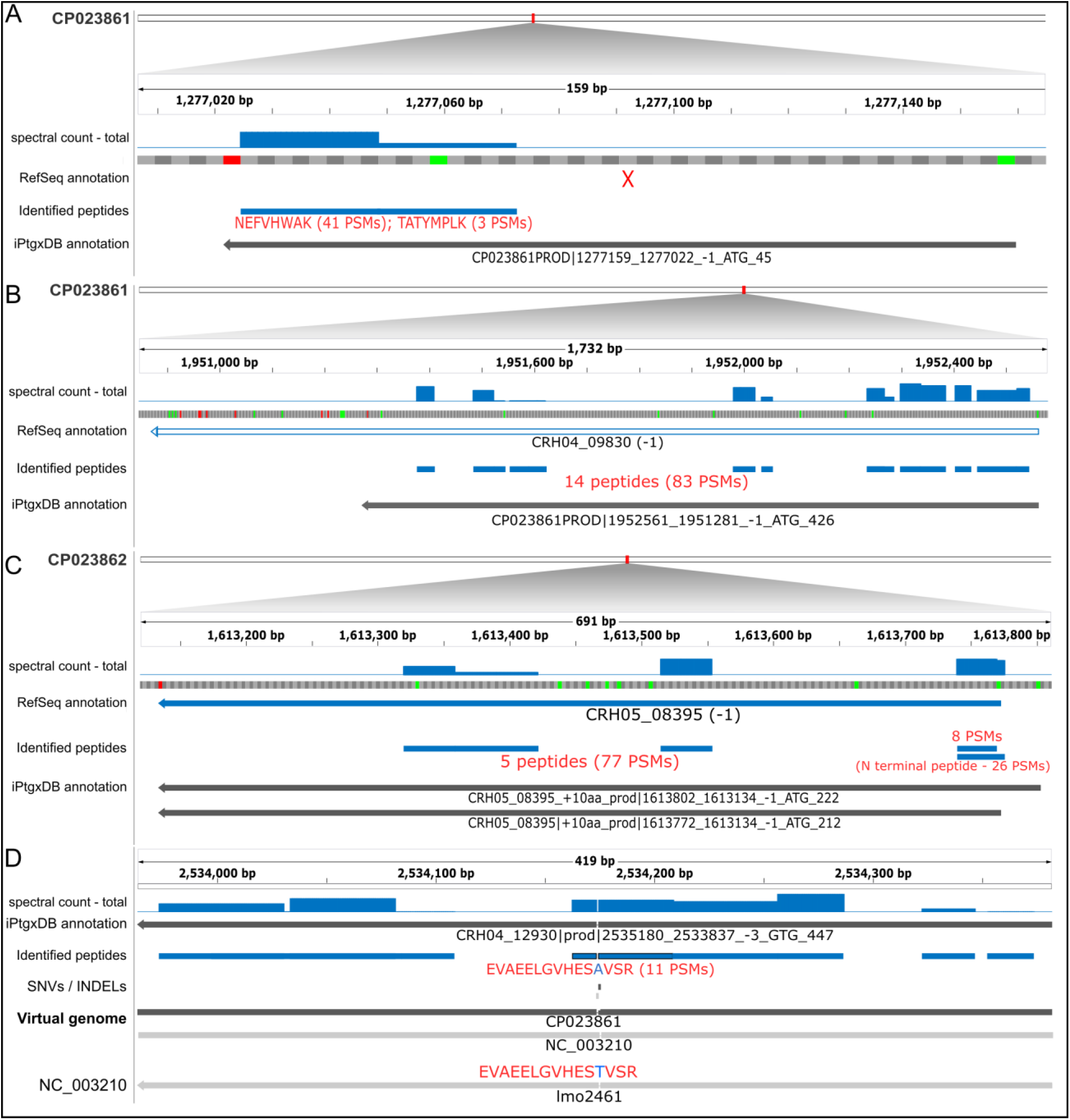
Peptide evidence for novelties identified by proteogenomics. Peptide evidence for **(A)** a new smORF, **(B)** an expressed pseudogene, **(C)** a new start site, and **(D)** a single amino acid variation (SAAV) uncovered in *L. monocytogenes* strain EGD-e and/or ScottA. Shown are respective genomic localizations. For A-C, the respective accession numbers of our assembly are given on the left above various annotation tracks. The iPtgxDB annotation is shown in dark grey, peptide evidence supporting novelties in blue, and a summary of these peptides and PSMs are shown in red. The sequences of the peptides implying the SAAV are shown in panel **(D)**. In the subfigure in panel **(D)**, two genome sequences are compared as a virtual genome, allowing us to overlay experimental evidence. All novelties were discovered by searching data against the strain-specific iPtgxDB. For simplicity, *in silico* predicted ORFs are not shown.

The iPtgxDBs are publicly available (https://iptgxdb.expasy.org/database/). These databases will support efforts in the *Listeria* community to find proteogenomic evidence for additional novel smORFs, as pioneered with the example of Prli42 [20]. This type of data has been instrumental in uncovering novelties in several model organisms including the eroded genome of an obligate plant symbiont [61]. Notably, the integration with global dRNA-seq data sets allowed identification of internal start sites of annotated proteins [62], similar to the N-terminomics study in *Listeria* [20].

### *L. monocytogenes* proteotype analysis reveals adaptation during stress

Overall, approximately 1,700 and 1,900 proteins were identified and quantified (for details see S8 Table) for EGD-e and ScottA, respectively, which represents a major advance for quantitative proteotype profiling studies in *Listeria*. S9 Table provides a summary over the respective library recovery percentage, data completeness and median CVs for both strains. Average correlation coefficients of biological replicates were above 0.98 indicating a very good reproducibility of sample analysis (data not shown). The reproducibility of the sample analysis was also reflected by the median CVs that ranged around 20% (S9 Table). Additionally, unsupervised hierarchical clustering revealed clustering of the biological replicates and a distinction across the different conditions tested (data not shown). Notably, the samples incubated with bile salts (duodenum-mimicking condition) demonstrated lower library recovery and a higher number of missing values. This was a result of the significantly lower amount of starting material due to decreased cell viability (Fig 3B). Fig 6 summarizes the proteins found to be differentially abundant in bile salts and low pH compared to the control condition for strains EGD-e (Fig 6A) and ScottA (Fig 6B), respectively. Very few proteins were differentially regulated under high osmolarity compared to the control (data not shown). The complete list of differentially regulated proteins under all conditions is included in S10 Table. In general, the condition mimicking those found in the stomach (low pH/pepsin) resulted in more quantitative changes compared to changes observed under high osmolarity for both strains (S10 Table). Also, more proteins were found to be differentially abundant upon perturbation in EGD-e compared to strain ScottA.

**Fig 6.**
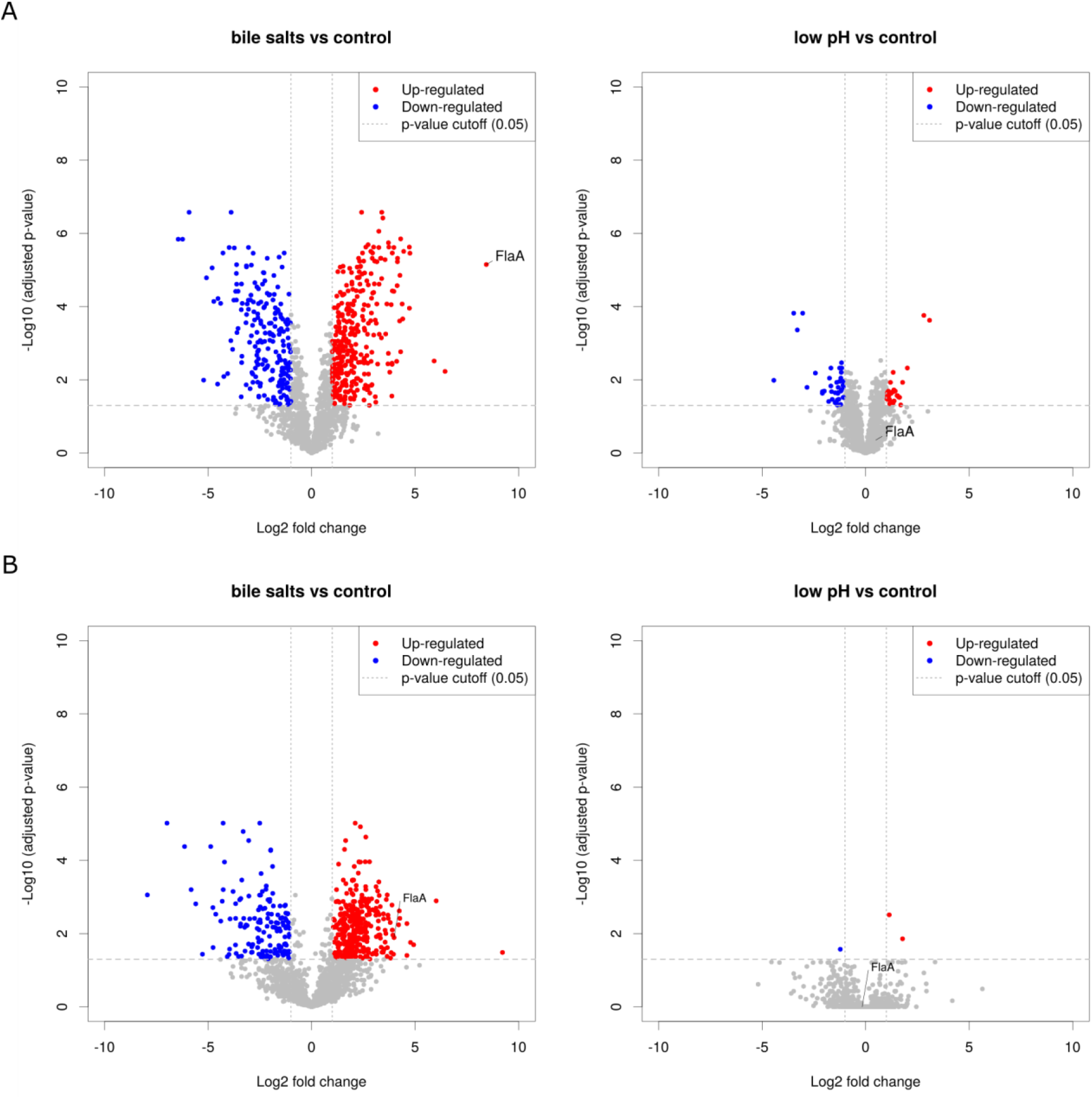
Differential protein abundance in EGD-e and ScottA cells upon exposure to conditions mimicking passage through the upper GI tract. Volcano plots depict the differentially abundant proteins for two conditions, i.e. stomach (low pH) and duodenum (bile salts) compared to control for **(A)** EGD-e and **(B)** ScottA. The adjusted p-value (multiple testing corrected, see Material and Methods) is shown as a horizontal gray dashed line, the log2 fold change cut-off of 1 as a vertical dashed gray line. Upregulated proteins are shown in red, downregulated proteins in blue; FlaA is labeled as one example.

Several studies have investigated the effect of bile salts on *L. monocytogenes* cells e.g. through gene knockout analyses, and several genes have been associated with resistance to bile (S11 Table). The majority of these proteins is present in our spectral libraries and one of them was specific to the EGD-e genome (CRH04_02340, which corresponds to the product of *pva*). Interestingly, at the proteotype level, only one of these proteins was found to be significantly differentially abundant when bacterial cells from both strains were incubated with bile salts compared to control condition (*bilEB;* EDG-e: CRH04_07370, ScottA: CRH05_07950; S11 Table). This likely indicates that the simplistic *in vitro* conditions developed only partially reflect the full complexity e.g. of the gallbladder fluid, which in other studies had been used from sacrificed animals [63].

One of the most highly differentially regulated proteins in both strains upon bile salt treatment was flagellin, the structural protein of *Listeria* flagella. This result was unexpected given that flagella are known to be thermoregulated and not formed in temperatures higher than 30 °C [64]. The gene encoding flagellin (*flaA*) is located right in between two predicted operons of flagella and motility associated genes ([65]; S11 Table. The majority of these genes was also detected at the protein level and are thus represented in the spectral libraries. For ScottA, the products of three additional flagellar hook-associated genes (*flgE*, CRH05_04145; *flgK*, CRH05_04185; *flgL*, CRH05_04190) were found to be upregulated upon incubation with bile salts. In contrast, in EGD-e, the products of the two component regulatory system (2CRS) *cheY* (CRH04_3620), a chemotaxis response regulator, and cheA (CRH04_03625) were upregulated in bile and low pH, or bile condition, respectively. Moreover, the list of differentially abundant proteins from all conditions was compared to the list of unique, strain-specific genes to investigate whether strain-specific alterations of the proteotype exist. For EGD-e, nine proteins encoded by the corresponding strain specific genes were differentially abundant across bile (7) and low pH (3) conditions with CRH04_001625 (a DUF1433 domain-containing protein) up in both (S10 Table). In contrast, fifteen ScottA strain-specific proteins were differentially regulated upon exposure to bile salts, including three transcriptional regulators (CRH05_01760, CHR05_02245, CRH05_11180; S10 Table). Together, these results suggest that different mechanisms are used by the strains in response to the bile salt stress.

### Flagella expression in bile salt condition could hint at a possible escape/survival mechanism

The most striking phenotypic change that was observed in both strains based on the DIA data (Fig 6) was the significant increase of flagellin protein FlaA, the structural protein of flagella, upon exposure to bile salts. The increased abundance of FlaA (Fig 6) was verified independently by parallel reaction monitoring mode (PRM) assays (Fig 7A). Additionally, as revealed by a PRM time-course experiment, protein levels of flagellin had increased after 15 min of incubation with bile salts (Fig 7B). A quantitative real-time PCR experiment was performed to assess *flaA* mRNA levels. Given that the FlaA protein levels had increased after 15 min of incubation in bile-containing medium, the qPCR was performed with bacterial cells grown in control and bile conditions for 5, 10, and 15 min. Strikingly, no significant upregulation of *flaA* mRNA was observed (Fig 7C). This suggests that protein production of FlaA is controlled at the post-transcriptional level.

**Fig 7.**
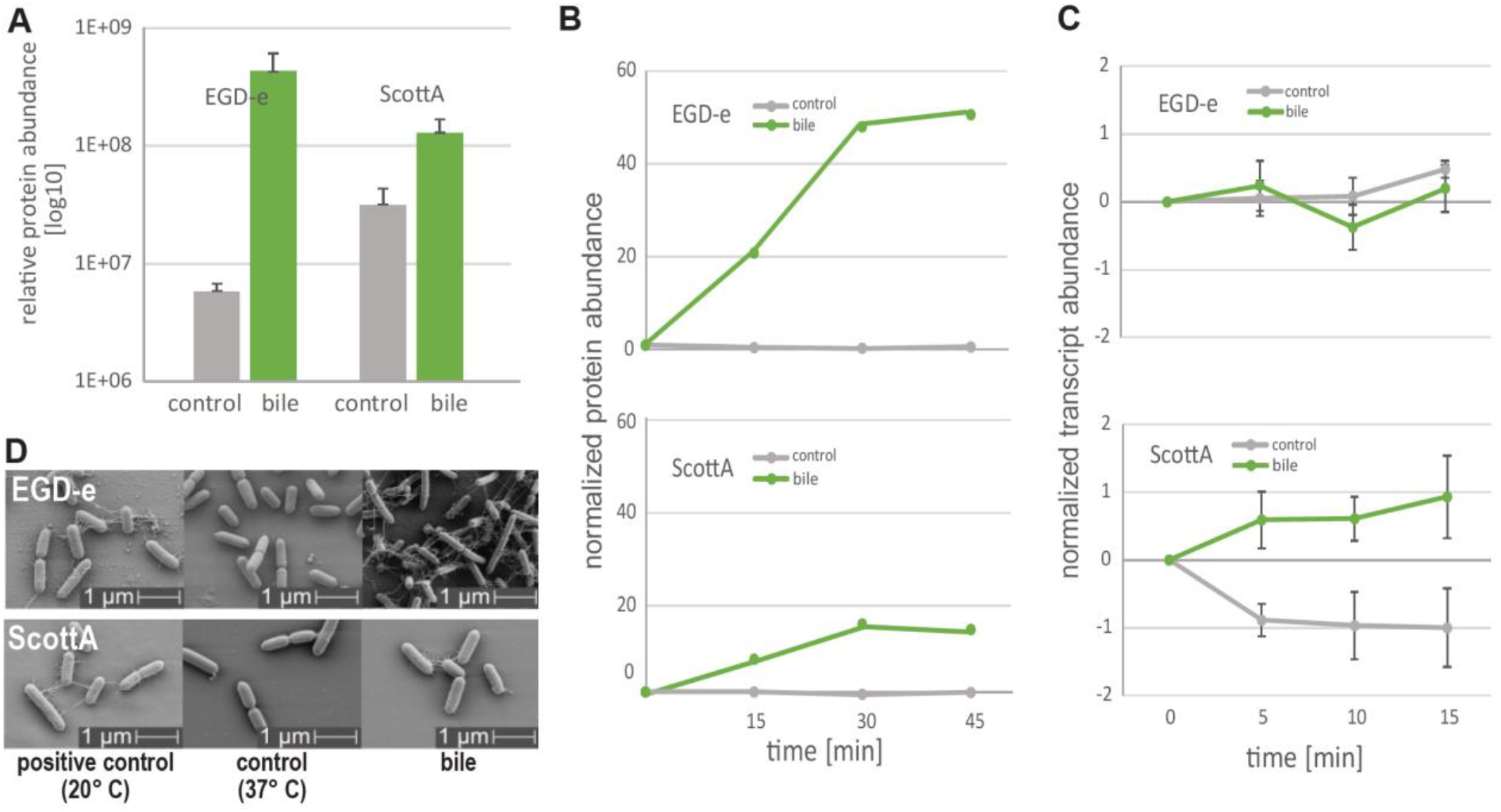
Production of flagella in *Listeria* strains incubated with bile salts at 37 °C. **(A)** Barplots of FlaA relative protein abundance based on PRM quantitation. **(B)** Protein and **(C)** RNA expression levels of *flaA* assessed in a time-course experiment using PRM and qPCR, respectively. **(D)** Representative scanning electron microscopy images of EGD-e and ScottA cells grown in indicated conditions.

To investigate whether the protein increase was the result of the formation of flagella, the cell surfaces of *L. monocytogenes* cells cultured in control and bile salt containing medium were stained and visualized using light microscopy. Cells grown in the presence of bile salts were flagellated to a larger extent than samples grown in control cultures (data not shown). In order to visualize the flagella at higher resolution, bacterial cells were analyzed by scanning electron microscopy (Fig 7D). The data confirmed that the flagella expression observed at 20 °C was highly reduced or absent at a temperature of 37 °C. However, upon exposure to bile salts, an increase in flagella expression was readily observed.

The above observations illustrate one valuable example of the additional unique information that can be obtained from quantitative next-gen proteomics but not from gene expression analysis. The presence of flagella under a bile salt stimulus could enable the bacteria to move away from that signal toward the intestinal mucus were they might subsequently lose their flagella and infect the host. Further experiments will be required to elucidate the function of flagella in the duodenum. The proteogenomic resources presented here will enable the *Listeria* community to add comprehensive quantitative proteotype profiling to their functional genomics technology set-up and uncover novel fascinating aspects of *Listeria* biology (or genotype-proteotype-phenotype relationships), some of which may be regulated at the proteotype level.

## Materials and Methods

### Bacterial strains and growth conditions

Bacterial strain EGD-e (serovar 1/2a) was derived from strain EGD, originally isolated from guinea pigs and used in studies of cell-mediated immunity [19], and differs quite substantially from EGD [65]. ScottA is a clinical strain (serovar 4b) that was isolated during the Massachusetts listeriosis outbreak in 1983 [44]. Cultures were grown to stationary phase in brain heart infusion (BHI) at 30 °C with shaking and then diluted 1:10 in BHI. Diluted cultures were incubated at 37 °C until the OD_600_ was 1. Cells were washed once with PBS and resuspended in the same volume of selected growth medium. Samples were incubated at 37 °C for 1 h with shaking, the medium was removed by centrifugation, and cell pellets were frozen until MS analysis. Three media were prepared to resemble conditions encountered in different parts of the upper GI tract [55] with buffered peptone water serving as control medium (BPW: 1% (w/v) peptone, 0.5% (w/v) NaCl, 0.35% (w/v) Na_2_HPO_4_, 0.15% (w/v) KH_2_PO_4_, pH 7.2). Low pH medium (stomach) was BPW, pH 4, 1000 units/ml pepsin. Bile salts medium (duodenum) was BPW, pH 7, 0.3% (w/v) bile salts. High osmolarity medium (jejunum) was BPW, pH 8, 0.3 M sucrose. To assess viability, seven serial dilutions of cells were prepared, and 10 μl of each sample were spotted on agar plates. Plates were incubated at 30 °C overnight, and colony counting was performed for the dilution where colonies were well separated from each other.

### Genome sequencing, assembly and annotation

Genomic DNA was prepared from overnight cultures of EGD-e and ScottA using the Sigma GenElute kit. An insert library was prepared, size selected with BluePippin (fragments > 10 kb), and sequenced on the PacBio RSII sequencing platform (1 SMRT cell per strain; P6-C4 chemistry). Initial pre-processing steps (read quality control, pre-assembly) and *de novo* genome assembly were carried out using HGAP3 [50] as described in detail before [43]. Subsequently, terminal repeats were trimmed, and the contigs were circularized and polished for two rounds using the PacBio pre-assembled reads. The EGD-e chromosome was aligned to the closest NCBI reference (GenBank accession: NC_003210) to adjust the start position according to the reference genome. For ScottA, the circularized contig was start aligned to the *dnaA* gene as described [53]. Both strains were also sequenced using Illumina MiSeq (2 × 300 bp paired end reads); raw fastq reads were mapped to the respective PacBio contigs using BWA-MEM v0.7.12 [66]. The final, high-quality genome sequences were submitted to NCBI GenBank and annotated with NCBI’s PGAP 4.0 [42].

### Functional annotation

In addition to the NCBI annotation, all protein coding genes were also annotated using Interproscan v5.30-69.0 [67] (restricted to hits with an e-value below 1e-5), adding information on Gene Ontology (GO) classification, protein domains, patterns, protein families, profiles, etc. Furthermore, we extracted functional annotations for the protein sequences using eggnog-mapper (v 1.0.3) [68], which transfers functional annotation from orthologous proteins present in EggNOG 4.5 [69]. Prophage sequences were identified and annotated using PHASTER; putative prophages have lower scores than those predicted to be intact [52]. A GO enrichment analysis of the proteins in the spectral library of each strain against all protein-coding genes of the respective strain was performed using the topGO package [70]. A Fisher’s exact test with a p-value cut-off of 0.01 was applied to identify significantly enriched GO categories across all three domains (i.e., BC, MF and CC) [71].

### Comparative genomics

A comparison of our two *de novo* assembled, complete genomes was carried out using Roary (v 3.7.0) [54] with standard parameters (minimum identity for blastp set to 50%, no paralog splitting). From the gene presence/absence output table, we extracted the core and strain-specific protein-coding gene clusters (S3 Table). The unique gene clusters and encoded proteins were subsequently used to identify the subset of proteotypic peptides that allowed quantification of the subset of proteins specific to each strain.

### Single-tube sample preparation for DIA/SWATH MS analysis

A 1-ml culture of approximately 10e^9^ bacteria was used as a starting material yielding roughly 100 μg of total protein. Cell pellets were reconstituted in 500 μl of 50 mM ammonium bicarbonate buffer, and 5 μg of phage endolysin Ply511 was added. Endolysin was produced in *E. coli* and purified by affinity chromatography as described earlier [72]. The amount and time of endolysin incubation was optimized to allow complete lysis (based on OD_600_ measurements, data not shown). Samples were incubated under fast end-to-end rotation for 15 min at 4 °C and then sonicated at maximum amplitude for 10 sec three times or until viscosity of water was reached. Samples were centrifuged for 10 min at maximum speed to remove debris, and a BCA protein assay was performed to assess the total protein concentration. For further preparation, 100 μg of total protein per sample were used. Proteins were denatured by heat (80 °C for 15 min) and addition of 0.1% acid-cleavable detergent. Proteins were reduced with 10 mM TCEP for 30 min at 25 °C and alkylated with 20 mM IAA for 30 min at 25 °C in the dark. Proteins were digested with LysC (1:300) for 3 h at 37 °C followed by trypsin digestion (1:100) for 16 h at 37 °C. Upon digestion, samples were acidified with 0.1% TFA and precipitated detergent was removed by centrifugation. Samples were desalted via mixed cation exchange chromatography and eluted in 1 ml 5% NH_4_OH/90% MetOH. Peptides were dried and reconstituted in 60 μl 5% ACN, 0.1% FA with addition of iRT standard (1:10 v/v).

### DIA/SWATH MS analysis

Peptides were analysed on an Orbitrap QExactive Plus mass spectrometer (Thermo Scientific) equipped with a nano-electrospray ion source (Thermo Scientific) and coupled to a nano-flow high pressure liquid chromatography (HPLC) pump with an autosampler (EASY-nLC II, Proxeon). Peptides were separated on a reversed-phase chromatography column (75-μm inner diameter PicoTip™ Emitter, New Objective) that was packed in-house with a C18 stationary phase (Reprosil Gold 120 C18 1.9 μm, Dr. Maisch). Peptides were loaded onto the column with 100% buffer A (99.9% H2O, 0.1% FA) at 800 bar and eluted at a constant flow rate of 200 nl/min with a gradient of buffer B (99.9% ACN, 0.1% FA) and a subsequent wash step with 90% buffer B. For the analysis of cell lysates, 3 μg of peptides were separated on a 50-cm heated column with a 120-min linear gradient of 5-35% B, followed by a 10 min gradient to 50% B and a 5 min gradient to 90% B. Between batches of runs, the column was cleaned with two steep consecutive gradients of ACN (10%-98%). The MS was operated in DDA mode, with an automatic switch between MS to MS/MS scans. High-resolution MS scans were acquired in the Orbitrap (120,000 resolution, automatic gain control target value 2×10^5^) within a mass range of 395 to 1500 m/z. The 20 most intense precursor ions (Top20) were fragmented using higher-energy collisional dissociation (HCD) to acquire MS/MS scans in the Orbitrap (30,000 resolution, intensity threshold 2.5×10^4^, target value 2×10^5^, isolation window 2 m/z). Dynamic exclusion was set to 30 s. Instrument performance was checked by regular quality control measurements using a yeast lysate and the iRT retention time peptide kit (Biognosys), both for DDA and DIA modes.

### Database search and spectral library construction

The RAW files were processed with Proteome Discoverer software, version 2.1 (http://planetorbitrap.com/proteome-discoverer) using the RefSeq protein databases of the *de novo* assembled EGD-e and ScottA strains with the iRT peptide sequences added. The processing workflow consisted of SequestHT [73] and Amanda [74] search nodes coupled with Percolator [75]. The following search parameters were used for protein identification: (i) peptide mass tolerance set to 10 ppm; (ii) MS/MS mass tolerance set to 0.02 Da; (iii) fully tryptic peptides with up to two missed cleavages were allowed; (iv) carbamidomethylation of cysteine was set as fixed modification, methionine oxidation and protein N-term acetylation were set as variable modifications. Percolator was set at max delta Cn 0.05, with target FDR strict 0.01 and target FDR relaxed 0.05. The spectral libraries were generated in Spectronaut v10 (Biognosys) using standard parameters including 0.01 peptide spectrum match (PSM) FDR and a Best N-filter with min three and max six fragment ions per peptide.

### DIA/SWATH MS sample acquisition and data analysis

HRM calibration peptides (Biognosys) were spiked into the DIA samples according to the manufacturer’s instructions. The samples were analyzed on the same LC-MS system as the DDA runs using identical LC parameters. The mass range m/z 375-1200 was divided into 20 variable windows based on density as described previously [76]. The MS was operated in DIA mode with an automatic switch between MS to MS/MS scans. High-resolution MS scans were acquired in the Orbitrap (35,000 resolution, automatic gain control target value 5×10^6^) within a mass range of 400 to 1220 m/z. DIA scans preceded an MS1 full scan in the Orbitrap (35,000 resolution, intensity threshold 3×10^6^) with a stepped NCE 22.5, 25, 27.5. Instrument performance was regularly checked as described above. All DIA data were analyzed directly in Spectronaut v10 (Biognosys) with standard settings (dynamic peak detection, automatic precision nonlinear iRT calibration, interference correction, and cross run normalization (total peak area enabled)). All results were filtered for a q-value of 0.01 (equal to an FDR of 1% on the peptide level). All other settings were set to default.

### Integrated proteogenomics search databases

IPtgxDBs were created for EGD-e and ScottA by combining the NCBI’s RefSeq protein annotation with Prodigal predictions, an *ab initio* gene predictor [60], and a modified six-frame translation [43]. Proteomics data from the DDA runs were searched against the iPtgxDB FASTA file of each strain individually using MS-GF+ v2017.01.13 [77] to identify evidence for novel ORFs, alternative protein start sites, and SAAVs compared to the reference genome sequence. The search was performed in target-decoy mode, with a precursor mass tolerance of 10 ppm, full-trypticity, maximum precursor charge of 4, carbamidomethylation of cysteine as fixed, and methionine oxidation, asparagine deamidation, and protein N-term carbamidomethylation as variable modifications. The search results were filtered for a PSM level FDR of 0.05%, which ensured an estimated protein level FDR below 1%. In addition, we assessed the proteotypicity of the identified peptides using an in-house version of the original PeptideClassifier [59], further extended to support proteogenomics in prokaryotes [43]. Only peptides that uniquely mapped to one protein (class 1a) were considered for protein identification. Moreover, following an earlier recommendation, we filtered the unambiguous peptides with an additional, variable PSM cut-off [78]: We required two PSMs/peptide for RefSeq annotated proteins; three PSMs per peptide for Prodigal predictions or N-terminal extensions to RefSeq proteins, and four PSMs per peptide for novel *in silico* predicted proteins. Data were overlaid on top of the GFF file and visualized in a genome browser.

### Statistical data evaluation

All proteomics experiments on EGD-e and ScottA were performed in biological triplicates, except for EGD-e bile, where we could only quantify duplicates. DIA mapping data were searched against the in-house generated spectral libraries using Spectronaut, and the list of quantified spectral features (fragment ions/peptide sequences) was retrieved. In MSstats3 (v3.12.2) [79], which is often used for downstream Spectronaut data processing, the features were log-transformed, and then subjected to median normalization. For feature summarization, the Tukey’s median polish algorithm was applied. Protein fold changes and their statistical significance were tested using at least five features per protein. Tests for significant changes in protein abundance across conditions were based on a family of linear mixed-effects models. P values were multiple testing corrected to control the experiment-wide FDR at a desired level using the Benjamini-Hochberg method. Proteins were considered differentially expressed if they showed a fold-change of 2 or higher and an adjusted p-value of 0.05 or lower.

### Validation via PRM MS

Peptides were separated by reversed-phase chromatography on a high-pressure liquid chromatography (HPLC) column (75-μm inner diameter, New Objective) that was packed in-house with a 15-cm stationary phase (ReproSil-Pur C18-AQ,1.9 micrometer) and connected to a nano-flow HPLC combined with an autosampler (EASY-nL1000). Peptides were loaded onto the column with 100% buffer A (99.9% H2O, 0.1% FA) and eluted at a constant flow rate of 300 nl/min with a 90 min stepped gradient from 3–25% buffer B (99.9% ACN, 0.1% FA) and 25-50%B. Mass spectra were acquired in PRM on an Orbitrap Fusion™ Tribrid™ Mass Spectrometer (Thermo Fisher Scientific). Spectra were acquired at 15,000 resolution (automatic gain control target value 5.0*10e^4^); peptide ions in the mass range of 340–1,400 were monitored. Stepped HCD collision energy was set to 27 (+/-5)%, maximum injection time to 22 ms. Monitored peptides and results were uploaded to PanoramaWeb (see Data access).

### Flagella staining for visualization under light microscopy

Flagella of *L. monocytogenes* were stained with Ryu stain using a wet-mount technique. Ryu stain was prepared before every experiment by mixing 1 part solution Il with 10 parts solution I. Solution I (mordant) was 10 ml of 5% aqueous solution of phenol, 2 g of tannic acid, and 10 ml of saturated aqueous solution of aluminum potassium sulfate-12 hydrate. Solution II (stain) was 12 g crystal violet in 100 ml of 95% ethanol. For staining, cells were grown in the desired conditions, and 3 μl of culture were transferred on a glass slide and covered with a cover slip leaving small air spaces around the edge. Slides were incubated 10 min at 25 °C for bacterial cells to adhere, and 10 μl of Ryu stain were applied at the edge of the cover slip. The stain was left to mix with the cell suspension by capillary action. Slides were incubated for 10 min at 25 °C and examined under the microscope at 100x (oil).

### Flagella visualization using scanning electron microscopy

Bacteria were cultured as described above, and 0.2 ml of suspension was applied for 20 min on 12-mm coverslips covered with 15 nm of carbon and coated with poly-L-lysine. After washing twice with PBS, cells were fixed with 2.5% glutaraldehyde in PBS at room temperature before immediately transferring the samples on ice for 2 h. Samples were then processed in a Pelco Biowave Pro+ tissue processor with use of microwave energy and vacuum. Briefly, the fixed samples underwent a second fixation step in 2.5% glutaraldehyde, before being washed and postfixed in 1% OsO4 followed by 1% uranyl acetate in water and dehydration by successive immersion in increasing concentrations of ethanol and finished by critical point drying out of ethanol. The dried coverslips were mounted on SEM aluminium stubs and sputter-coated with 5 nm of platinum/palladium. SE-images were recorded at 2 kV in a Zeiss Gemini 1530 FEG.

### Nucleic acid extraction, purification, and cDNA synthesis

Bacterial nucleic acids were extracted using a phenol-chloroform protocol adapted from a previous report [80]. Briefly, the bead-beating step was carried out with 500 μl of 0.1-mm Zirconia/Silica Beads in a Mixer Mill MM301. Lysis was performed in two rounds of 4 min each with 5 min rest on ice. Nucleic acids were precipitated by addition of 0.1 volume 3 M sodium acetate and 0.6 volume ice-cold isopropanol. The pellet was resuspended in RNase/DNase-free water and subsequently purified with the AllPrep DNA/RNA kit (Qiagen) according to the manufacturer’s instructions. DNA isolated from the control culture grown in BHI at 37 °C served as positive control. DNA and RNA were quantified with the Qubit® Fluorometer 3.0. RNA was isolated from all samples, and 0.8 μg were used for cDNA synthesis (in triplicate) with the TaqMan® Reverse Transcription Reagents kit according to the manufacturer’s instructions using the random hexamer technique. Some reverse-transcription reactions were carried out without enzyme as a control for the absence of DNA in the samples.

### qPCR

The reaction was performed in a final volume of 10 μl containing 1X SYBR Green Mix, 0.5 μM each of the *flaA* primers (forward: 5’-GCTGGTCTTGCAGTTGTTACTCGTATG-3’; reverse: 5’-CTAATTGACGCATACGTTGCAAGATTG-3’) and 1 μl of the diluted DNA template using the SYBRTM Select Master Mix and run on a Rotor-Gene 6000 PCR system with the following program: 50 °C for 2 min, 95 °C for 2 min, followed by 40 cycles at 95 °C for 15 min. Three biological replicates were analyzed with samples analyzed in duplicate by qPCR. Raw data were processed with the LinRegPCR program to determine the Cq values, and Python 3.7.0 was used for analysis and to create plots.

## Author contributions

A.R.V. and M.P.P performed all experiments except those noted below. V.G. performed qPCR. S.G. performed PRM experiments. A.R.V., M.P.P., S.G. and V.G. analyzed data. Y.S. and M.J.L. supervised V.G. and Listeria experiments in the Loessner laboratory. M.P.P and B.W. conceived the project, C.H.A conceived genomics and proteogenomics aspects. A.R.V, M.P.P, C.H.A. and B.W. designed research. C.H.A., M.P.P, A.R.V and B.W. wrote and revised the manuscript. All authors commented and provided feedback on the manuscript.

## Acknowledgements

The authors thank Ulrich Omasits for bioinformatic support in the early stage of the project and Michael Schmid (both Agroscope) for contributions to the *de novo* genome assemblies. We would like to thank Patrick Studer (ETH Zurich) who helped with culture of the *Listeria* strains and J. R. Wyatt for text editing. BW acknowledges support from the Swiss National Science Foundation (SNSF) under grant 31003A_160259, CHA acknowledges support from the SNSF for ARV under grant 31003A-156320.

## Data access

The genome sequences for ScottA and EGD-e are available from NCBI Genbank under accession numbers CP023862 and CP023861, respectively. IPtgxDBs for both strains are available at https://iptgxdb.expasy.org. Targeted MS experiments can be accessed via Panorama (https://panoramaweb.org/660xuF.url). The proteomic data that were used as a basis to develop the spectral libraries and the DIA measurements are available from MASSIVE under ftp://MSV000083881@massive.ucsd.edu (reviewer access login: MSV000083881/ PWD: Listeria_2019). Selected datasets will also be accessible from the Listeriomics web server.

## Supporting information

**S1 Table. Bacterial strains.**

**S2 Table. Overview of genome properties of strains EGD-e and ScottA.**

**S3 Table. Overview of core genes and genes specific to ScottA and to EGD-e.** The subsets of core and strain-specific protein-coding genes are listed for both strains (see separate Excel table). Detailed functional annotation is provided. The 14 genes that were missing in the NCBI reference sequence of ScottA (CM001159.1) are highlighted in orange. Strain-specific genes that are represented in the spectral libraries are marked “TRUE” in the column “present in spectral_library”.

**S4 Table. Master table.** The list of all protein-coding genes in the *de novo* assembled strains, their functional annotations (NCBI and additional functional annotations; see Materials and Methods), proteomics expression evidence obtained using MS-GF+ after filtering for PSM level FDR of 0.05% (protein level FDR 1%), as well as the results of a reciprocal best BLAST hit analyses with the ListiList strain EGD-e (facilitating integration with other datasets) and its corresponding Uniprot accession are shown in a separate Excel file.

**S5 Table. Summary of annotation clusters in the iPtgxDBs.**

**S6 Table. Proteomics evidence for the novelties identified through the proteogenomics search of DDA data for *L. monocytogenes* strains EGD-e and ScottA.** See separate Excel table.

**S7 Fig. An incorrectly predicted pseudogene in ScottA.** Protein evidence for a 207-amino acid protein (3 peptides, 17 PSMs) predicted by Prodigal in *L. monocytogenes* strain ScottA. It starts from an alternative start codon TTG, which codes for leucine in frame −1; the corresponding longer RefSeq protein (282 amino acids) is wrongly predicted as a pseudogene. Both proteins are annotated as phosphosugar binding transcriptional regulators, and both harbor the RpiR-like SIS (sugar isomerase) protein domains (IPR035472).

**S8 Table: Summary information of precursors, peptides, and protein groups identified for EGD-e and ScottA over all conditions using DIA.**

**S9 Table. Summary of library recovery percentage, data completeness, and median CVs for the DIA dataset.**

**S10 Table. List of differentially abundant proteins.** Proteins identified as differentially abundant in the bile, low pH, and high osmolarity conditions for both strains. Strain-specific genes are highlighted in orange. See separate Excel table.

**S11 Table. Candidate genes for bile resistance and operons for flagellar genes.** List of *Listeria* genes previously shown to be involved in bile resistance and genes that are part of two operons with flagellar-related and motility genes that flank the *flaA* gene (Bécavin C, et al. Comparison of widely used Listeria monocytogenes strains EGD, 10403S, and EGD-e highlights genomic variations underlying differences in pathogenicity. MBio. 2014;5: e00969–14). See separate Excel table.

