## Supporting information for "A proteogenomic resource enabling integrated analysis of *Listeria* genotype-proteotype-phenotype relationships"

S6 Table. Proteomics evidence for the novelties identified through the proteogenomics search of DDA data for *L. monocytogenes* strains EGD-e and ScottA. 5

**S1 Table. Bacterial strains.**

| Strain no. | Name | Isolate type | Serotype |
|---|---|---|---|
| NF-L101 | EGD-e | Virulent laboratory strain (guinea pig)* | 1/2a |
| NF-L725 | ScottA | Virulent laboratory strain from a clinical isolate (foodborne)** | 4b |

Source of strains: * Mackaness GB. The immunological basis of acquired cellular resistance. J Exp Med. 1964; 120:105-120. Glaser P, et al. Comparative genomics of *Listeria* species. Science. 2001;294: 849–852. ** Fleming DW, et al. Pasteurized milk as a vehicle of infection in an outbreak of listeriosis. N Engl J Med. 1985;312: 404-407. Briers Y, et al. Genome sequence of *Listeria monocytogenes* ScottA, a clinical isolate from a food-borne listeriosis outbreak. J Bacteriol. 2011;193: 4284–4285.

**S2 Table. Overview of genome properties of strains EGD-e and ScottA.**

| | ***L. monocytogenes* EGD-e** | ***L. monocytogenes* ScottA** |
|---|---|---|
| Genbank accession # | CP023861 | CP023862 |
| # Chromosomes | 1 | 1 |
| Size of the chromosome (bp) | 2,944,523 | 3,030,813 |
| G+C content (%) | 37.9 | 37.9 |
| Coverage (PacBio) | 260 | 283 |
| Coverage (Illumina MiSeq) | 302 | 435 |
| Total number of protein-coding genes | 2,887 | 2,979 |
| Number of rRNA operons (16S-23S-5S) | 6 | 6 |
| Number of tRNA genes | 67 | 67 |
| Number of pseudogenes | 29 | 30 |
| Prophages | 2 (probable) | 2 (intact) + 1 (probable) |

**S3 Table. Overview of core genes and genes specific to ScottA and to EGD-e.**

The subsets of core and strain-specific protein-coding genes are listed for both strains (see separate Excel table). Detailed functional annotation is provided. The 14 genes that were missing in the NCBI reference sequence of ScottA (CM001159.1) are highlighted in orange. Strain-specific genes that are represented in the spectral libraries are marked "TRUE" in the column "present in spectral_library".

**S4 Table. Master table.**

The list of all protein-coding genes in the *de novo* assembled strains, their functional annotations (NCBI and additional functional annotations; see Materials and Methods), proteomics expression evidence obtained using MS-GF+ after filtering for PSM level FDR of 0.05% (protein level FDR 1%), as well as the results of a reciprocal best BLAST hit analyses with the ListiList strain EGD-e (facilitating integration with other datasets) and its corresponding Uniprot accession are shown in a separate Excel file.

**S5 Table. Summary of annotation clusters in the iPtgxDBs.**

A. EGD-e (65,393 proteins)

| Annotation source (prefix) | # total CDS | # total clusters | # total new clusters | # total new reductions | # total new extensions | # total clusters | # total ids |
|---|---|---|---|---|---|---|---|
| RefSeq (refseq) | 2,919 | 2,919 | 2,919 | 0 | 0 | 2,919 | 2,919 |
| Prodigal (prod) | 2,877 | 2,877 | 34 | 80 | 84 | 2,953 | 3,117 |
| *In silico* ORFs (orf) | 67,980 | 48,406 | 45,466 | 31 | 18,847 | 48,419 | 67,461* |

*Excluded were 1,989 proteins smaller than 6 amino acids; 29 entries were all-pseudo annotated and excluded from the iPtgxDB; 50 shorter entries had indistinguishable internal start sites and were also excluded from the iPtgxDB.

B. ScottA (67,150 proteins)

| Annotation source (prefix) | # total CDS | # total clusters | # total new clusters | # total new reductions | # total new extensions | # total clusters | # total ids |
|---|---|---|---|---|---|---|---|
| RefSeq (refseq) | 3,010 | 3,010 | 3,010 | 0 | 0 | 3,010 | 3,010 |
| Prodigal (prod) | 2,970 | 2,970 | 48 | 75 | 89 | 3,058 | 3,222 |
| *In silico* ORFs (orf) | 69,931 | 49,726 | 46,677 | 23 | 19,433 | 49,735 | 69,355* |

*Excluded were 2,122 proteins smaller than 6 amino acids; 30 entries were all-pseudo annotated and excluded from the iPtgxDB; 53 shorter entries had indistinguishable internal start sites and were also excluded from the iPtgxDB.

**S6 Table. Proteomics evidence for the novelties identified through the proteogenomics search of DDA data for *L. monocytogenes* strains EGD-e and ScottA.** See separate Excel table.

**S7 Fig. An incorrectly predicted pseudogene in ScottA.** Protein evidence for a 207-amino acid protein (3 peptides, 17 PSMs) predicted by Prodigal in *L. monocytogenes* strain ScottA. It starts from an alternative start codon TTG, which codes for leucine in frame -1; the corresponding longer RefSeq protein (282 amino acids) is wrongly predicted as a pseudogene. Both proteins are annotated as phosphosugar binding transcriptional regulators, and both harbor the RpiR-like SIS (sugar isomerase) protein domains (IPR035472).

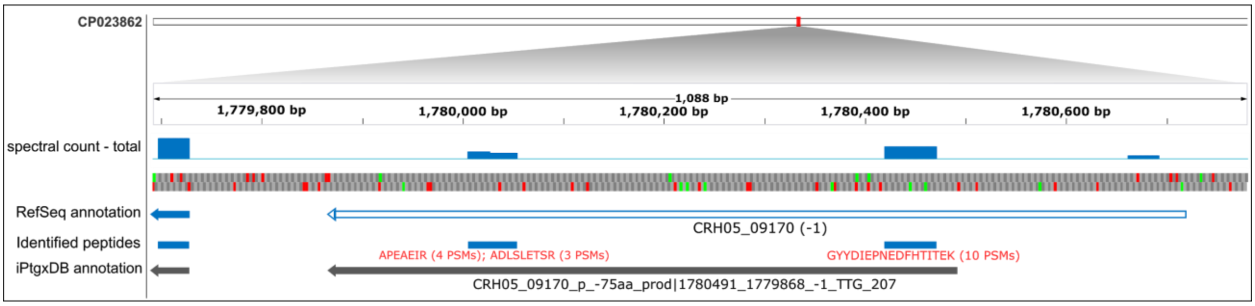

**S8 Table: Summary information of precursors, peptides, and protein groups identified for EGD-e and ScottA over all conditions using DIA.**

| | EGD-e | ScottA |
|---|---|---|
| **Precursors** | 22,993 | 25,585 |
| **Peptides** | 16,725 | 16,635 |
| **Modified Peptides** | 19,002 | 21,169 |
| **Protein Groups** | 1,708 | 1,876 |
| **Precursor recovery from library [%]** | 90.8 | 97.5 |

**S9 Table. Summary of library recovery percentage, data completeness, and median CVs for the DIA dataset.** Part of this dataset is also shown in Fig 6.

| | **EGD-e** | | | | **ScottA** | | | |
|---|---|---|---|---|---|---|---|---|
| **Condition** | **Control** | **Low pH** | **High osmolarity** | **Bile salts** | **Control** | **Low pH** | **High osmolarity** | **Bile salts** |
| **Data completeness** | 85% | 84% | 81% | 35% | 91% | 80% | 87% | 51% |
| **Library recovery** | 85% | 85% | 82% | 36% | 95% | 88% | 92% | 65% |
| **Median CVs** | 17% | 19% | 17% | 20% | 20% | 21% | 17% | 81% |

**S10 Table. List of differentially abundant proteins.** Proteins identified as differentially abundant in the bile, low pH, and high osmolarity conditions for both strains. Strain-specific genes are highlighted in orange. See separate Excel table.

**S11 Table. Candidate genes for bile resistance and operons for flagellar genes.** List of *Listeria* genes previously shown to be involved in bile resistance and genes that are part of two operons with flagellar-related and motility genes that flank the *flaA* gene (Bécavin C, et al. Comparison of widely used *Listeria monocytogenes* strains EGD, 10403S, and EGD-e highlights genomic variations underlying differences in pathogenicity. MBio. 2014;5: e00969–14). See separate Excel table.
